# Sweep Dynamics (SD) plots: Computational identification of selective sweeps to monitor the adaptation of influenza A viruses

**DOI:** 10.1101/110528

**Authors:** Thorsten R. Klingen, Susanne Reimering, Jens Loers, Kyra Mooren, Frank Klawonn, Thomas Krey, Gülsah Gabriel, Alice C. McHardy

## Abstract

Monitoring changes in influenza A virus genomes is crucial to understand its rapid evolution and adaptation to changing conditions e.g. establishment within novel host species. Selective sweeps represent a rapid mode of adaptation and are typically observed in human influenza A viruses. We describe Sweep Dynamics (SD) plots, a computational method combining phylogenetic algorithms with statistical techniques to characterize the molecular adaptation of rapidly evolving viruses from longitudinal sequence data. To our knowledge, it is the first method that identifies selective sweeps, the time periods in which these occurred and associated changes providing a selective advantage to the virus. We studied the past genome-wide adaptation of the 2009 pandemic H1N1 influenza A (pH1N1) and seasonal H3N2 influenza A (sH3N2) viruses. The pH1N1 influenza virus showed simultaneous amino acid changes in various proteins, particularly in seasons of high pH1N1 activity. Partially, these changes resulted in functional alterations facilitating sustained human-to-human transmission. In the evolution of sH3N2 influenza viruses, we detected changes characterizing vaccine strains, which were occasionally revealed in selective sweeps one season prior to the WHO recommendation. Taken together, SD plots allow monitoring and characterizing the adaptive evolution of influenza A viruses by identifying selective sweeps and their associated signatures.

## Introduction

Influenza A viruses are rapidly evolving pathogens causing respiratory infections with high morbidity and mortality in the human population^1^. Annual influenza epidemics result in 3 to 5 million reported infections and up to 250,000–500,000 cases of death^1^. Currently, the viral subtypes sH3N2 and pH1N1 are circulating in the human population. The H3N2 virus was introduced into the human population in 1968 and is endemic ever since. The swine-origin H1N1 subtype emerged in the 2009 influenza pandemic and was subsequently referred to as the 2009 pH1N1 virus. It replaced the formerly circulating seasonal H1N1 subtype^2,3^. The negative-sense RNA genome consists of eight segments that encode for 14 viral proteins^4^. A constant arms-race between the human immune system and the virus results in continuous adaptation of the viral genome. These changes facilitate the virus to escape the host’s immune response elicited through vaccination or previous influenza infections^5^. Alterations in the major glycoproteins − hemagglutinin (HA) and neuraminidase (NA) − and genomic reassortment that change the viral antigenicity are defined as antigenic drift and result in re-occurring epidemics of seasonal influenza viruses^6^. Continuous antigenic changes of circulating strains require a re-evaluation of antigenically predominant strains by the WHO twice a year at the end of each season, leading to a recommendation for the vaccine composition for the following year^7,8^. The establishment of an antigenically new virus strain into an immunologically naive human population causes pandemics, mostly due to alterations in the receptor binding protein HA^9^.

Measuring the impact of natural selection plays a crucial role in molecular evolution, as it determines the genomic constitution and diversity of a population^10^. Directional evolutionary processes boost the fitness of individuals by introducing advantageous amino acids. The respective alleles rise in frequency in the viral population, consequentially reducing variation. This process is called a ‘selective sweep’^11^. Hereafter, we use the term ‘sweep-related change’ when referring to an amino acid exchange rising in frequency due to a selective sweep. Due to linkage within genomic segments, the advantageous amino acid exchange promoting the sweep and other changes in close genomic proximity jointly rise in frequency, leading to an increase of linkage disequilibrium (i.e. nonrandom associations between genomic regions). Thus, selective sweeps are of profound interest since they represent a rapid shift of the whole genotype carrying the selected amino acid change. The increasing number of available viral genomes that are provided by modern sequencing techniques allows us to conduct a genome-wide analysis for recent selective sweeps to gain insight into the within-host evolution and adaptation of human influenza A viruses.

Several methods were developed to detect and measure the effect of directional evolution for viral populations. We previously described Allele Dynamics (AD) plots, which characterize the evolutionary dynamics of sets of amino acid changes, indicating those most likely to be under positive selection using population level time-series data sets of genetic sequences^12^. Similarly, nextflu.org provides a web-based visualization of changes in allele or clade frequencies in the HA protein of circulating seasonal influenza viruses^13^. Related work from *Luksza and Lassig*^14^ builds upon clade frequencies to forecast influenza lineages with acquired fitness advantages in the viral population. Other methods to analyze natural selection are based on non-synonymous to synonymous mutation rates (*dN/dS*)^15^. Synonymous changes are assumed to be neutral, while a relative excess of non-synonymous changes, i.e. *dN/dS* > 1, indicates positive selection^15^. *dN/dS* is either calculated by counting synonymous and non-synonymous substitutions^16,17^ or by estimation using maximum likelihood models^16,18^. This statistic is not applicable to detect selective sweeps, as a large number of synonymous substitutions could occur after a substitution rose to fixation in a selective sweep, which would result into a *dN/dS* ratio smaller than 1 and the conclusion that this site is not evolving under positive selection^11^. Moreover, *dN/dS* was originally developed for the analysis of divergent species and the interpretation of *dN/dS* > 1 as positive selection may not be correct for single populations^19^. To elude the limitations of *dN/dS*, *Bhatt et al.* developed a statistic based on site frequencies to calculate site- or segment-specific adaptation rates^20^. Furthermore, several other approaches have been developed to identify selective sweeps using accelerated substitution rates^11^, a skew in the allele frequency spectrum^18^ or an excess of linkage disequilibrium^21^. Detecting the season or the time period in which a specific amino acid change involved in a selective sweep emerged, would improve comparative molecular studies of natural selection and indicate changes with potential effect on the viral fitness. This could aid to uncover the drivers of adaptive evolution in viral populations.

Here, we describe Sweep Dynamics (SD) plots, which allow analyzing the population-level phylodynamics of influenza virus proteins or proteins of other rapidly evolving organisms from longitudinal samples of genetic sequences. A statistical evaluation reveals selective sweeps, and in addition the season in which they occurred and the associated individual amino acid changes. We used the SD plots for a genome-wide characterization of directional selection in pH1N1 influenza viruses since their introduction into the human host. In all proteins under consideration, we inferred sweep-related changes that indicate human-adaptive changes after its emergence in 2009, several of which were in structural proximity to known mammalian host adaptation sites. Furthermore we detected sweep-related changes in antigenicity- and avidity-changing sites of the sH3N2 influenza virus hemagglutinin that correlate with newly emerging antigenic variants in the human population and show the value of SD plots for vaccine strain selection.

## Results

### Sweep Dynamic (SD) Plots

Sweep Dynamics (SD) plots are an extension of the AD plot technique that we previously described^12^. SD plots analyze the evolutionary dynamics of alleles, representing individual amino acid changes within the viral population. The dynamics of reconstructed amino acid changes (alleles) in an homogeneous, constant-sized viral population can be described by a Fisher model^22,23^. This model is also considered valid with changing population sizes^24-26^ and seasonal influenza virus populations have been modeled as homogenous due to their rapid spread around the globe^19^. Genetic drift and selection are acting on alleles, resulting in changes of allele frequencies (ratio of copies of one genetic variant relative to the population size)^27,28^. In this population, an allele under directional selection rises faster in frequency than alleles without a selective advantage. Amino acid changes (alleles) that increase in frequency swiftly over time are thus more likely to be under directional selection than other alleles with lower frequencies. We apply this criterion in the SD plots method together with a statistical evaluation, to pinpoint those amino acid changes that increase significantly faster in frequency than others and thus might provide a selective advantage. Changes are inferred under consideration of the evolutionary relationships, to consider their relatedness and avoid overcounting changes resulting from denser sampling of specific parts of the viral population.

From the sequences for a particular protein from a viral population sample, a phylogenetic tree is inferred and amino acid changes in its evolution are reconstructed that map to the individual branches in the tree (Material & Methods). The frequencies of circulating isolates that have inherited a particular amino acid change are deduced from the phylogenetic tree by counting the number of viral isolates descending from the branch where a particular amino acid change was introduced. This is done separately for each season (Figure 1A&B). Other than the AD-plots, which combine amino acid changes at different positions of the coding sequence into alleles when they share a branch or occur in close proximity in the phylogeny, SD plots analyze individual amino acid changes for their potential effect on viral fitness. The visualization of SD plots highlights these identified sweep-related changes and their variation in frequency over time. Specifically, the following procedure is carried out:

**Figure 1:**
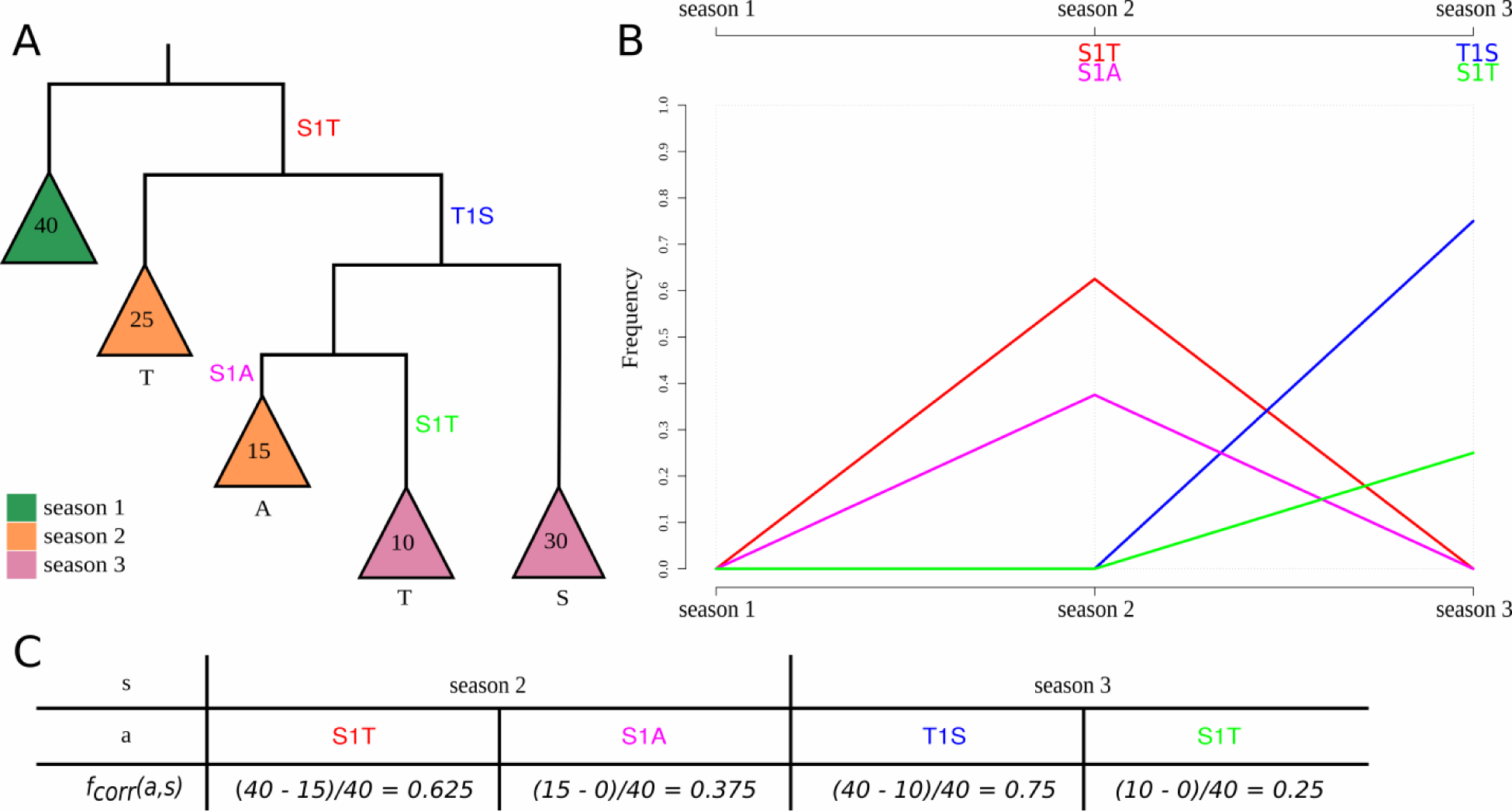
Phylogeny demonstrating the frequency correction, the corresponding SD plot and frequencies of amino acid changes. This is a detailed illustration of how the frequency correction handles individual amino acid changes that occur at the same position. The tree *T* **(A)** consists of five main subtrees sampled from three seasons (colored in green, orange and purple). Each season *s* exhibits forty isolates |*L_T,s_*| = 40 ∀ *s*). *A_T_* contains four amino acid changes that occur on different internal branches: S1T (red), T1S (blue), S1A (pink) and S1T (light green). We consider S1T (red) and S1T (light green) as two separate amino acid changes with individual frequencies as they occur on different branches of the tree. Consequently, both are calculated separately and occur in the SD plot **(B)**. Amino acid changes S1T (red) and S1A (pink) rise in frequency in the second season and disappear in the last season while T1S (blue) and S1T (light green) arise in the third season in the corresponding SD plot. Note that the frequencies for change S1T (red) and for T1S (blue) are corrected while the frequencies for S1A (pink) and S1T (light green) do not require a correction **(C).**

Using a phylogenetic tree *T* and a set of reconstructed amino acid changes *A_T_* for all internal branches, the frequency of isolates in the population that carry a specific amino acid change 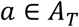 is calculated as follows: let *L_T_* represent the set of viral isolates assigned to the leaves in *T*, with each viral isolate 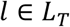 being labeled with a time stamp *s* indicating the season in which it was sampled and *L_T_, s* representing the subset of sequences of *L_T_* labeled with time stamp s. Starting from the root, we perform a level order traversal, visiting each edge *e* in the tree *T*, and calculate the frequency for each individual amino acid change 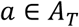 that is located on the edge *e* as follows: Let *T_a_* be the subtree that is rooted at the node with the in-edge *e*. Let *A_T_a__* be the set of amino acid changes in the subtree *T_a_*, with *A_T_a__* ⊂ *A_T_* and 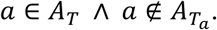. The frequency *f*(*a, s*) of an individual amino acid change *a* in season *s* is defined as: 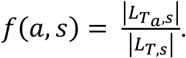 It represents the ratio of all isolates in the subtree that have acquired the amino acid change *a* within season *s* relative to the number of all isolates within the designated season *s*.

As we consider individual positions, the prevalence of one amino acid change per position is affected by the emergence of a more recent amino acid change at the same position, making it necessary to adjust the frequency *f*(*a, s*) at position *p* in the alignment. The frequency *f*(*a, s*) of *a* occurring at position *p* is adjusted when another amino acid change *β* occurs at *p* in the subtree *T_a_* − a process referred to as frequency correction, as follows: let there be *n* amino acid changes at *p* in the subtree *T_a_*, i.e. |*A_T_a__*| = *n*. Each subtree *τ_β_* is rooted at the node with the in-edge that represents the amino acid change 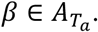. The set 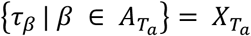 represents the *n* subtrees that are contained in the tree *T_a_*. Let 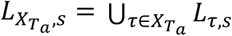 be the set of all leaves with the time stamp in all subtrees in *X_T_a__*. Note that 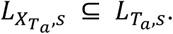 The corrected frequency *f_corr_*(*a, s*) of amino acid change *a* in season *s* is then defined as: *f_corr_*(*a, s*) = 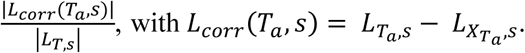 The frequency of an amino acid change *a* is adjusted by excluding isolates *L_X_T_a__, s_* that are subject to a more recent substitution (Figure 1). The SD plots always apply *f_corr_*(*a, s*) because either the frequency needs to be corrected or whenever no additional amino acid change occurs in the phylogenetic tree *T* after the amino acid change *α*, the term 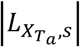 is zero and automatically results in the uncorrected case (*f*(*a, s*)). Thus, the equation *f_corr_*(*a, s*) ≤ *f*(*a, s*) holds. Note that this allows an individual analysis of the same amino acid exchange introduced several times in the phylogeny on different branches (e.g. S1T (red & light green) in Figure 1) and their distinct evolutionary trajectories.

To identify a selective sweep and the associated amino acid change, we define two criteria: an amino acid change should show a significant increase in frequency relative to the previous season. For each season, we only report changes that were not reported as significant before. For each amino acid change, we test the null hypothesis that the number of viruses in the viral population carrying this change is equal or lower than in the previous season. We evaluate the significance of frequency changes for each amino acid change *a* using |*L_corr_*(*T_a_, s*) | and |*L_corr_*(*T_a_*, *s* + 1)| Fisher’s exact test^29^, using and over consecutive seasons *s* and *s* + 1. A significant *p*-value (*p* ≤ 0.05) indicates that an amino acid change significantly increased in frequency in the current season relative to the previous one. To correct for multiple testing, we adjusted the *p*-values with the Benjamini-Hochberg procedure controlling the false discovery rate at level *α* = 0.05^30^.

For each dataset, the seasons from the second to the last one are tested, comparing each season to the preceding one. We report amino acid changes for the season *s* of their first predominant occurrence (*f_corr_*(*a, s*) > 0.5; it occurs in more than 50% of the isolates within the designated time period) as in *S*teinbrück and McHardy^12^ in combination with a significant *p*-value (*p* ≤ 0.05).

The results are visualized in SD plots (Figure 2 & 3). These provide a detailed overview of the emergence of individual amino acid changes in the viral population and in which season a sweep took place. The changes in frequency of sweep-related changes are depicted as trajectories over consecutive seasons. The season of their first predominant occurrence is indicated with an asterisk. In each season, the amino acid exchanges of all sweep-related alterations are listed in a panel above the graph. Within the panel, they are bottom-up ordered with ascending frequency.

**Figure 2:**
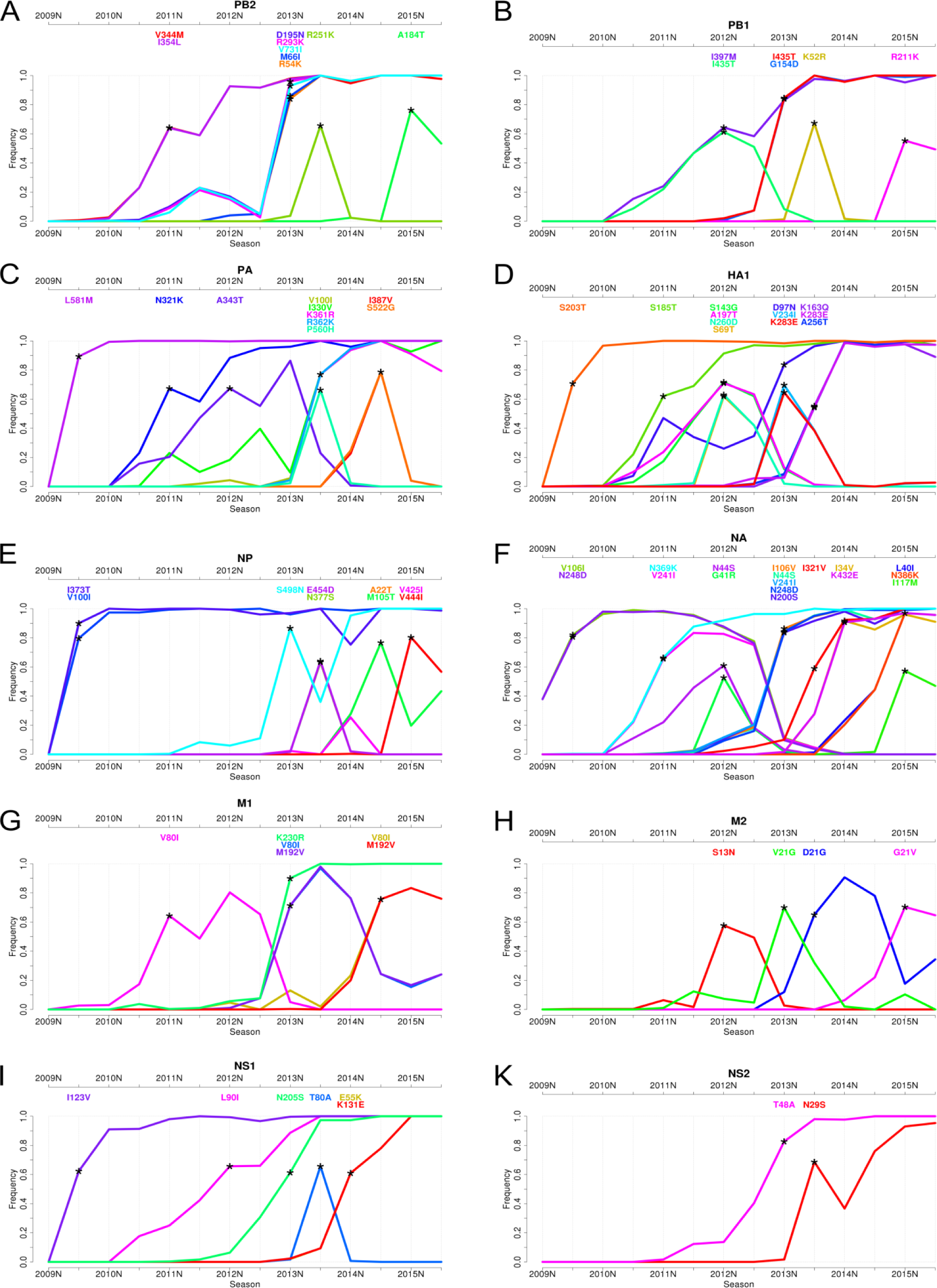
Sweep Dynamics (SD) plots for ten pandemic H1N1 influenza A proteins. These plots show the alterations in frequency of sweep-related changes over consecutive time periods in the proteins PB2, PB1, PA, HA, NP, NA, M1, M2, NS1 and NS2. They are ordered from **(A)** – **(K)** based on decreasing segment size on which they are encoded. Each plot shows the seasons (from 2009S to 2015S) on the x-axis and the frequency on the y-axis. The initial emergence of a sweep-related amino acid change is indicated by an asterisk pinpointing the first season in which it significantly rises in frequency to a frequency of more than 50%. Sweep-related changes are listed in a panel above the graph and are identically color coded as the curve in the plot. Within the panel, they are bottom-up ordered with ascending frequency.

**Figure 3:**
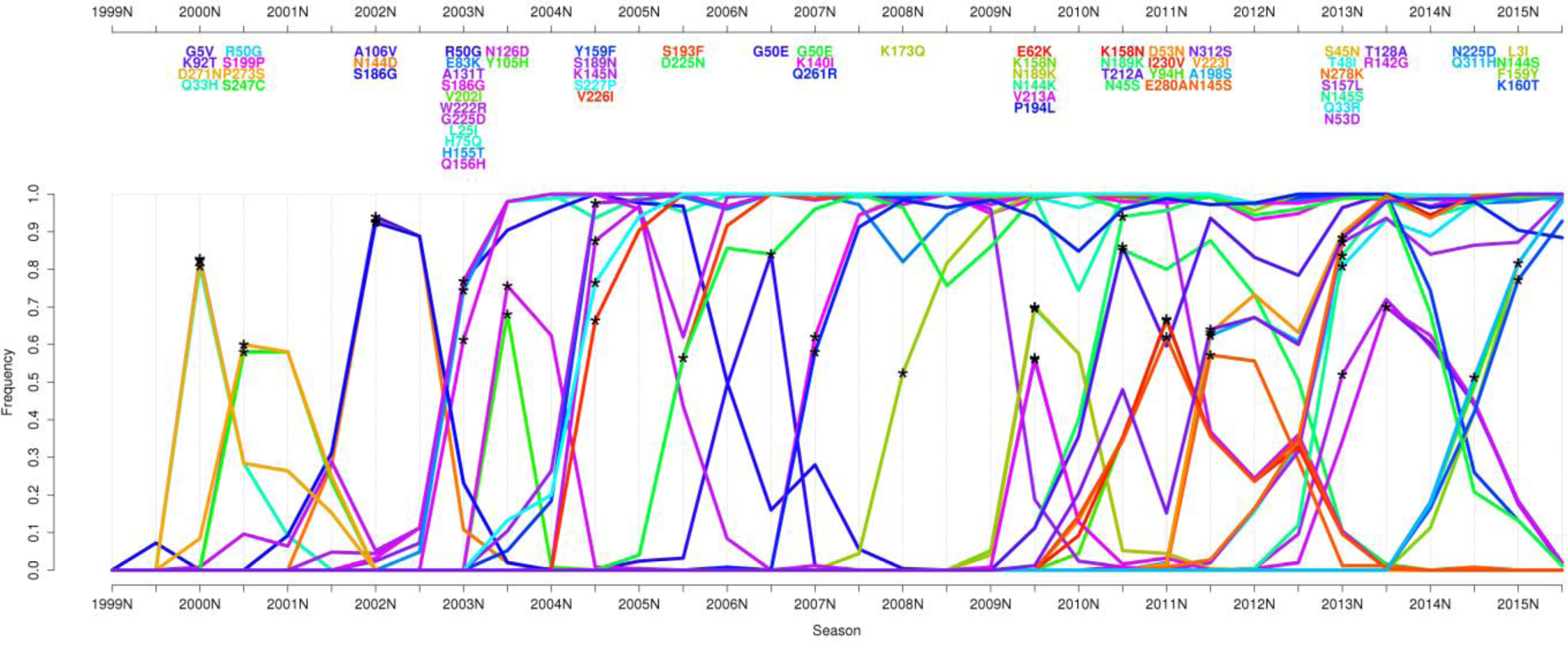
Sweep Dynamics (SD) plot for the seasonal H3N2 influenza A hemagglutinin. This plot shows alterations in frequency of sweep-related changes over consecutive time periods for the influenza A/H3N2 hemagglutinin. It depicts the seasons (from 1999S to 2015S) on the x-axis and the frequency on the y-axis. Sweep-related changes are listed in a panel above the graph and are identically color coded as the curve in the plot. Within the panel, they are bottom-up ordered with ascending frequency.

### Selective sweeps in the evolution of pH1N1 influenza viruses

We applied SD plots to detect selective sweeps in the past evolution of pH1N1 influenza viruses. We analyzed nucleotide and amino acid sequences of ten proteins (HA, NA, M1, M2, NS1, NS2, NP, PA, PB1 and PB2) collected since the appearance of the virus in the beginning of 2009 until the end of September 2015^2^. Sequences were assigned to influenza seasons using the common definitions for seasons in the Northern and Southern hemisphere (Data & Methods). For pH1N1 influenza, the data covered fourteen seasons from 2009N to 2015S (Figure 2, Supplementary Table 1). To investigate the structural relationships of sweep-related changes in the proteins, we mapped sweep-related changes in HA1, NA, NP and the polymerase (PB1, PB2 and PA) onto the respective structures (Material & Methods).

The SD plots analysis indicated selective sweeps and associated changes for the 2009S, 2011N, 2012N, 2013N, 2013S, 2014N, 2014S and 2015N seasons. The most sweep-related changes (twelve and seventeen, respectively) were detected for the surface proteins HA and NA and the fewest were found for NS2 (two changes) over all thirteen seasons. Newly arising changes often occurred simultaneously, i.e. were detected in the same season, with up to five changes detected in a protein in one season (2013S in PA and NA, 2013N in PB2), and with changes occurring simultaneously in multiple proteins (Figure 2). In addition to changes providing a selective advantage, some of these are likely hitchhikers without notable effect on fitness. With hitchhikers, we refer to (almost) neutral changes that are introduced into a sequence shortly before or after a change causing a selective sweep. Hitchhikers then rise in frequency together with beneficial changes due to genomic linkage and are thus detected as being sweep-related. Technically, the simultaneous occurrence of amino acid changes on the same branch prevents a computational distinction between their potential effects, as they are ancestral to the same set of leaf nodes. Besides linkage within a genomic segment, there is also a strong linkage across all influenza segments^31-33^, resulting in hitchhikers from other segments being carried along to higher frequencies by functionally relevant changes. The visualization of evolutionary dynamics in the SD plots can identify some of these ‘sweep-related hitchhikers’ as those decreasing in frequency after their emergence and not becoming fixed.

For the 2009S season, we detected a selective sweep at seven amino acid positions throughout the viral genome (in PA (L581M), HA (S203T) (position 202 in *Otte et al.*^34^), NP (V100I, I373T), NA (V106I, N248D) and NS1 (I123V)) (Figure 2C, 2D, 2E, 2F and 2I, Supplementary Table 1), which were also described in *Otte et al.*^34^. All of these changes rapidly became predominant in this season and continued to rise in frequency until they were close to fixation in 2010N, suggesting that the new amino acid was present in nearly all sampled isolates in this season. These findings are in agreement with the study of *Elderfield et al.*, in which 2009S was identified as the time of the first wave of pH1N1 activity after its emergence^35^. The pandemic waves were defined based on large viral prevalence with a phase of low viral prevalence in between. Five of seven detected sweep-related changes were also described in this study as emerging changes in the first wave of the pandemic, with frequencies close to 100% in the second wave in 2010N, similar to the frequencies we observed in our analysis (frequencies between 91% and 99%, with slight differences most likely due to differing time frames and datasets in both analyses). With the exception of the V106I and N248D changes in NA, all sweep-related changes detected in 2009S were still fixed in the population in 2015. The frequency of the amino acid change V106I in NA decreased from 2011S onwards, as it was replaced by the I106V sweep-related change in 2013N. The frequency of V106I accordingly dropped rapidly from 78% to 11% between 2012S and 2013N and continued to decrease until it disappeared in 2014S. From 2011S onwards, the frequency of N248D also decreased to 0% in 2014S. This amino acid change re-emerged in 2013N and replaced the strain with the first change. This repeated emergence indicates a particular relevance of this change regarding viral fitness. All seven sweep-related changes detected in 2009S have been reported to induce functional changes that might have facilitated viral adaptation to the human population: we previously described that all changes with the exception of NS I123V mediated increased pH1N1 influenza virus pathogenicity in a mouse model^36^. Moreover, all seven adaptive mutations were associated with enhanced respiratory droplet transmission in a ferret model, suggesting their crucial role to sustain with the mammalian host^34^. The ability of influenza A viruses to transmit from human-to-human is considered to be key in pandemic spread. Hereby, particularly the receptor binding properties of HA play an important role. We have also previously shown that HA S203T increases binding to α2,6-linked sialic acids (corresponds to position 202 in *Otte et al.*^34^) that are predominantly expressed in the upper respiratory tract of humans, a key site in virus transmission among humans^34,37,38^.

Both residues I100 and T373 in NP are solvent exposed (*RSA* = 0.32 and = 0.26, respectively); T373 is in close proximity to N319 (10.2 Å measured from Cα to Cα, Figure 5A), at which an N to K change promotes adaptation to mammalian cells^39^. Elevated virulence was also found for substitutions V106I in NA and I123V in NS1^40^. The V106I and N248D substitutions enhance viral stability at low pH, which confers replicative fitness and likely promotes virus spread^41^. Notably, through the N248D change, the protein acquires a negative charge (at neutral pH) that is solvent-exposed (*RSA* = 0.56) and located within < 12 Å vicinity to residues belonging to the three-dimensional structure of the active center of NA^42,43^ (Figure 5B).

**Figure 4:**
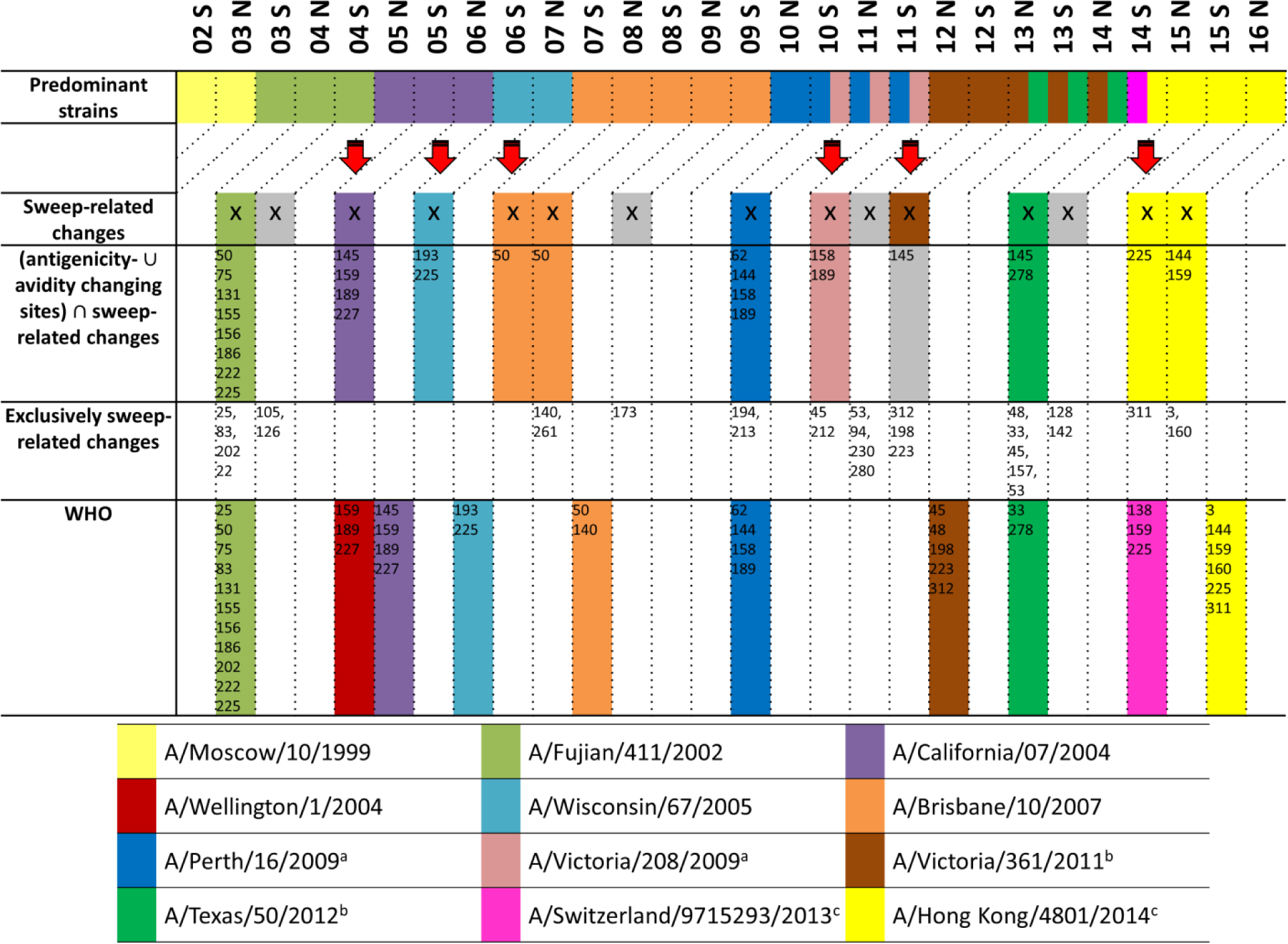
Comparison of predominant seasonal H3N2 influenza A strains, SD plots results and recommendations made by the WHO. From 1999N until 2015S, eleven antigenically different sH3N2 influenza strains were selected for production of the seasonal influenza vaccine, named Moscow/10/1999, A/Fujian/411/2002, A/Wellington/1/2004, A/California/7/2004, A/Wisconsin/67/2005, A/Brisbane/10/2007, A/Perth/16/2009, A/Victoria/361/2011, A/Texas/50/2012, A/Switzerland/9715293/2013 and A/Hong Kong/4801/2014^2,75-83^, which are indicated by colors as labeled in the legend (first row). For the SD plots analysis, seasons are marked (with an X in the second row) if sweep-related changes distinguish the respective vaccine strain from the previous one. False positive or false negative results are marked in grey. The third row lists sweep-related changes that also are antigenicity or avidity changing sites, while the fourth row shows detected sweep-related sites that are neither known to change the avidity or the antigenicity. The selection of a vaccine strain takes place two seasons before the vaccine is available. Any prediction of newly arising antigenically novel strains should therefore not be compared to the current predominant strain, but to the predominant strain two seasons after the detection. This comparison of the WHO selections and the SD plots results with the predominant strain two seasons later is indicated by the diagonal lines in the upper part of the plot (fifth row). ^a^The strain A/Victoria/208/2009 was predominant in 10S, 11N and 11S but was antigenically indistinguishable from A/Perth/16/2009^3,84,85^. ^b^The strains A/Victoria/361/2011 and A/Texas/50/2012 were predominant in 13N, 13S and 14N but A/Texas/50/2012 was antigenically more effective^81,86,87^. ^c^In 14S, two different clades containing the strains A/Switzerland/9715293/2013 and A/Hong Kong/4801/2014, respectively, were rising in frequency at the same time and antigenically differed from the previous vaccine strain A/Texas/50/2012^82^.

**Figure 5:**
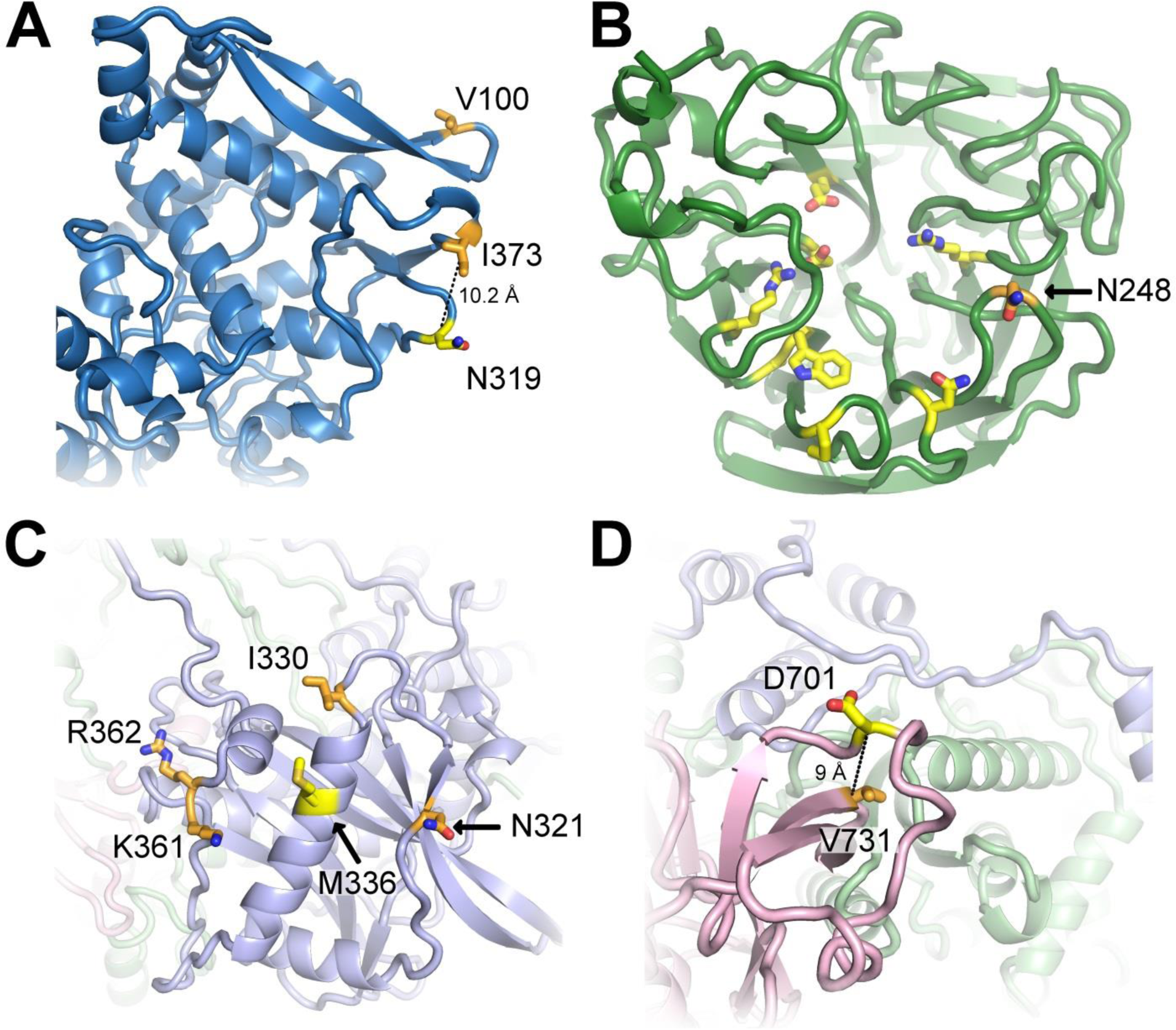
Representation of sweep-related changes on protein structures. We mapped individual sweep-related changes on the corresponding models shown as cartoon with interesting residues shown as sticks and colored by atom-type (yellow and orange for carbon, red for oxygen and blue for nitrogen). In **(A)**, the structure of NP is shown in blue with sweep-related mutated residues T373 and I100 colored orange and mammalian-adaptation site N319 yellow. The distance between C?-atoms of T373 and N319 is 10.2 Å (dotted line). The structure in **(B)** depicts NA (green) with residue N248 (orange) in proximity (<12 Å) to the active center (yellow)^41^. In **(C)** and **(D)** we show views on the structural model of the polymerase with subunits PB2 (pink), PB1 (light green) and PA (light blue). Sweep-related changes at residues N321, I330, K361 and R362 (orange) in PA cluster around M336 (yellow) that increases polymerase activity in mammalian cells **(C)**^45^. The structure of polymerase subunit PB2 (pink) is shown in **(D)** and highlights the sweep-related mutation V731 (orange), which is in close vicinity (≤9 Å) to position 701 that enhances mammalian host adaptation^39,48^.

We detected a second selective sweep in season 2011N, corresponding to the third wave of the pandemic^35^. We identified seven accompanying changes in PB2, PA, HA, NA and M1 (Figure 2A, 2C, 2D, 2F and 2G, Supplementary Table 1). The changes V244M and I354L (PB2), N321K (PA), S185T (HA) and N369K (NA) came close to fixation in later seasons, with frequencies of around 90% in 2012N. Notably, their rise in frequency was slower in comparison to the changes from 2009S, which reached a frequency of nearly 100% only one season after they were detected. The change S185T in HA was reported to enhance receptor binding avidity^44^, while N321K in PA increased polymerase activity^35^. This residue is exposed on the structure (*RSA* = 0.41) and close to M336 (Figure 5C), which was reported to foster adaptation by increasing polymerase activity in mammalian cells^45^. Furthermore, the amino acid changes N369K and V241I in NA were described as permissive mutations which enhanced viral fitness in oseltamivir resistant viruses^46^. As the amino acid change V241I decreased in frequency in 2013N (Figure 2F) and is buried in the protein (*RSA* = 0), this could indicate that site N396K (exposed, *RSA* = 0.1; uncharged to positively charged) might be the key amino acid change that confers antiviral resistance.

After the third global wave in 2011N, the pH1N1 influenza virus continued to circulate in the human population with lower activity throughout 2012 in a wide range of countries, especially in Europe and Eastern Asia^47^. The SD plots indicated a selective sweep in 2012N accompanied by eleven changes in the proteins PB1, PA, HA, NA, M2 and NS1 (Figure 2B, 2C, 2D, 2F, 2H and 2I, Supplementary Table 1). Eight of these changes (I435T in PB1 and all changes in HA, NA and M2) decreased in frequency right after this selective sweep and disappeared completely in 2013S. The A343T change in PA also was lost a year later in 2014S. This behavior is striking in comparison to the previously detected sweeps in 2009S and 2011N, where a majority of sweep-related changes became fixed (except V241I (NA) and V80I (M1)). For the selective sweep in 2012N, only the I397M amino acid change in PB1 and the L90I amino acid change in NS1 continued to be present in the circulating viral population and became fixed in 2013S (Figure 2B and 2I), suggesting their relevance for the sweep while the remaining changes could have been hitchhikers.

In 2013N, the pH1N1 influenza virus re-emerged with more activity in the countries where activity was low in 2012^47^. Here the SD plots revealed twenty-two sweep-related changes in all proteins, except for PA (Figure 2, Supplementary Table 1). PB2 and NA had the most changes, with five amino acid changes each rising rapidly in frequency and coming close to fixation within one season. Seventeen of these twenty-two amino acid changes were still fixed at the end of 2015S, suggesting that some of these changes provide a selective advantage. The sweep-related change V731I (PB2) is located in the vicinity (9 Å) of residue 701 (Figure 5D), a position known to be involved in mammalian host adaptation^39,48^. Further sweep-related changes not rising to fixation occurred in HA (V234I and K283E), M1 (V80I and M192V) and M2 (V21G).

We detected a sweep in 2013S with sixteen sweep-related changes in nine proteins, with six changes in PB2 (R251K), PB1 (K52R), PA (P560H), NP (E454D and N377S) and NS1 (T80A) decreasing quickly in frequency and showing frequencies close to 0% one season after their emergence. The other sweep-related changes were still fixed in the viral population in 2015S and thus could be of functional relevance, including V100I, I330V, K361R and R362K in PA, K163Q, K283E and A256T in HA and I321V in NA (Figure 2C, 2D and 2F). Especially the changes I330V, K361R and R362K in line with N321K (season 2011N) cluster significantly around the mammalian-adaptation site 336 on the structure of PA that increases polymerase activity (Figure 5C)^49^. This might suggest that these PA changes might also affect viral polymerase activity, as a hallmark of influenza disease severity in humans. We tested for enrichment of fixed sweep-related sites within a radius of < 13 Å in the vicinity of site 336 (hypergeometric distribution; *N* = 716, *K* = 31, *n* = 6, *k* = 4; *H*_0_: fixed sweep-related changes occur with the same probability in vicinity to site 336 and in the remaining protein region; *P* = 4 × 10^−5^; N being all sites in the protein model, K the median number of sites in a radius of 13 Å around a site in the protein model, n all detected sweep-related changes which remained fixed, k the number of fixed sweep related changes in the cluster). The change K283E in HA (site 283 is exposed; *RSA* = 0.41) results in a change of charge from positive to negative and is located in the stalk region that plays a key role in the induction of neutralizing antibodies^50^.

The 2014N and 2014S seasons showed sweep-related changes only for a small number of proteins. In 2014N, two changes were detected in both NA and NS1 (Figure 2F, 2I), which all rose to fixation and still circulated in 2015S. In 2014S, changes in the proteins PA, NP and M1 were identified subsequently. While I387V and S522G in PA disappeared in 2015S, M105T and A22T in NP were maintained at frequencies of around 40% (Supplementary Table 1). The V80I and M192V changes in M1 were already detected in 2013N, but disappeared due to back mutations and re-emerged, indicative of a functional relevance.

Similar to 2013S, many sweep-related changes detected in 2015N decreased in frequency in 2015S. Exceptions are L40I and N386K in NA, which remained close to fixation in 2015N and seem most likely to provide a selective advantage. Six sweep-related changes in PB2 (A184T), PB1 (R211K), NA (I117M), NP (V425I, V444I), M2 (G21V) decreased in frequency but still remained predominant, with frequencies around 50% or higher in 2015S (Figure 2A, 2B, 2D, 2E and 2H, Supplementary Table 1).

### Selective sweeps in the evolution of sH3N2 influenza

To investigate the past evolution of the seasonal sH3N2 influenza virus, we analyzed sequence data of the 34 seasons from 1999N to 2015S. The sH3N2 influenza virus has been circulating in the human population since 1968, but before 1999N sequence data only few data are available. In the SD plots analysis we therefore focused on more recent years, ensuring a sufficient data coverage for each season (Figure 3, Supplementary Table 2).

Overall, we identified fifteen seasons in the analyzed time period with sweep-related changes in the HA protein. For each detected selective sweep, we investigated whether it indicated the emergence of an antigenically novel strain that became predominant by matching the sweep-related changes to the amino acid changes that were reported for the viral strain by the WHO. If not stated directly for the predominant strain, the changes reported for the subclade including the predominant strain were used for matching to the WHO reported strains, as they distinguish a novel viral variant from previous ones. We further assessed the coherence between detected selective sweeps and novel antigenic variants by comparing the sweep-related sites to known antigenicity- and avidity-changing sites. We recently reported twenty-three antigenicity-altering sites organized in five patches on the structure of HA playing a role in the past antigenic evolution of sH3N2 viruses^51^, a subset of which together with position 193 were also reported by *Koel et al.*^52^. Of the sites influencing the receptor binding properties, i.e. avidity changing, we focused on the sites 193, 222 and 225^53^, as well as 145^54^, which is also an antigenicity-altering site^51^, bringing the set of considered sites to twenty-six.

Of the fifteen seasons in which sweep-related changes were detected in the HA protein, eleven included changes at these twenty-six sites (Figure 4). Of the eleven sweeps with relevant changes, three matched and indicated the emergence of a novel antigenic variant one season before the strain became predominant (A/Fujian/411/2002 strain, A/California/7/2004 strain and A/Perth/16/2009 strain) (Figure 4). In two cases, a newly emerging strain was matched and detected two seasons before predominance (A/Wisconsin/67/2005 and A/Brisbane/10/2007) (Figure 4). This is ideal for vaccine strain selection, as new strains to be included in the vaccine formulation are decided on one year in advance. Three cases matched and indicated a newly emerging strain in the season in which it was predominant (A/Victoria/208/2009, A/Texas/50/2012 and A/Hong Kong/4801/2014). Interestingly, for two strains (A/Brisbane/10/2007 and A/Hong Kong/4801/2014) we identified the associated sweep-related changes over two consecutive seasons. Only the appearance of the A/Victoria/361/2011 strain coincided with a sweep without changes at known antigenicity or avidity sites (Figure 4), which indicates that our list of antigenicity or avidity altering sites likely is not complete. The four sweeps without changes in antigenically- and avidity-altering sites did not include changes observed for antigenically novel predominant strains reported by the WHO.

In comparison to the strain recommendations by the WHO, sweeps predicted with the SD plots and including changes at antigenicity- or avidity altering sites identified newly emerging strains at the same time in three cases (e.g. A/Fujian/411/2002) or up to two seasons before in six cases (e. g. A/Brisbane/10/2007; Figure 4). Notably, the SD plots analysis did not identify sweeps corresponding to A/Wellington/1/2004 and A/Switzerland/9715293/2013, which were false positive vaccine strain recommendations that did not match the dominantly circulating strain^55-57^. In some cases practical issues such as the lack of a vaccine strain with sufficient growth properties in eggs, which might have prevented recommendation of a suitable vaccine strain candidate by the WHO. SD plot analysis indicated suitable strains in the majority of cases using a fixed computational procedure, and therefore seems suitable for supporting the vaccine strain decisions of the WHO experts. Notably, different from the established WHO procedure, no interpretation of hemagglutination inhibition assay data was required for this result. This could be an additional benefit, as recently, HI assays fail to agglutinate many circulating sH3N2 viruses^58^.

We further applied the SD plots to sequences collected prior to the WHO meeting of the respective seasons. In this scenario we produced a genealogy and accompanying SD plot for each season, thus excluding data from later time points also from tree inference. Except for minor differences in sweep-site detection and occasionally altered frequencies, this did not alter the suggested updates of the vaccine strains obtained when analyzing a tree generated across the entire time period (Figure 4). The plots are provided at https://github.com/hzi-bifo/SDplots.

## Discussion

We describe a new technique called SD plots that combines phylogenetics with a statistical analysis to detect selective sweeps and associated individual amino acid changes under directional selection from longitudinal samples of population-level sequence data. Due to the rapid evolution of influenza viruses, backmutations or repeated mutations at individual sites are common in certain protein regions, requiring elucidation of the evolutionary histories of the proteins for statistical analysis, to allow consideration of such effects. A unique aspect of the method is that it suggests individual amino acid changes under directional selection, instead of sites, which are candidates for experimental studies.

For pH1N1 viruses, the SD plots identified sweep-related changes that clustered on the protein structure together with known host adaptation sites, which could be studied experimentally for their relevance in host adaptation. This was the case for instance in the polymerase proteins, where such “adaptation patches” included sites with changes known to increase polymerase activity in mammalian cells and to promote adaptation to the human host since the establishment of pH1N1 in 2009. Future wet-lab experiments could focus on position 731 in the pH1N1 PB2 protein (a potential novel adaptation site), positions 330, 361 and 362 in pH1N1 PA and positions 54, 66 in pH1N1 PB2 (potentially promoting polymerase activity). Most sweep-related changes identified in HA of sH3N2 viruses occurred at sites of known relevance for immune evasion. SD plots in combination with filtering by antigenic patch sites and avidity changing sites enabled the timely detection of newly emerging antigenic variants for the sH3N2 viruses. Using these as suggested updates of the vaccine strain resulted in a better match to future predominant strains than those used (also in a retrospective testing scenario, where only isolates with a collection date prior to the respective vaccine selection meeting were used). An ideal set up for vaccine strain prediction would consider only those sequences sampled *and sequenced* until to the vaccine selection meeting of the WHO, e.g. from GISAID (http://platform.gisaid.org). However, there is a larger time gap between isolation and submission to GISAID, indicating that the respective labs do not submit all the data they provide to the WHO directly also to public repositories, and preventing an entirely realistic evaluation. For example, there were thirteen sequences with isolation and submission date prior to the vaccine selection meeting in 2008S available and sixty-three sequence with isolation date prior to 2008S but with a submission date from season 2009N.

Notably, SD plots could also be used to study the evolutionary dynamics of other rapidly evolving populations from longitudinal sequence samples, such as clinical intra-patient HIV data.

## Data & Methods

### Data download and sampling

Protein and nucleotide coding sequences were downloaded from the NCBI flu database^59^. For both the pH1N1 influenza and sH3N2 influenza subtype, we downloaded sequences of the proteins HA, NA, M1, M2, NS1, NS2, NP, PA, PB1 and PB2. The pH1N1 influenza virus was analyzed from the emergence of the virus in early 2009 until the end of September 2015, while sH3N2 influenza was studied from October 1998 until the end of September 2015. We downloaded sequences with the full date available, to properly assign sequences to influenza seasons. We used the standardized definition for the Northern hemisphere season (N) that begins on 1^st^ October of the previous year and ends on 31^st^ March and the Southern hemisphere season (S) that begins on 1^st^ April and ends on 30^th^ September in the same year, as before^12^.

We limited our analysis to isolates represented by both a nucleotide and a corresponding amino acid sequence. To account for variable numbers of sequences per season, we sampled the data with replacement, generating the same number of sequences per season for our analysis (300 sequences for pH1N1, 250 sequences for sH3N2). For HA, the amino acid numbering based on the mature protein without the signal peptides was used; corresponding to seventeen and sixteen amino acids for pH1N1 influenza viruses and sH3N2 influenza viruses, respectively^60,61^. In an additional experiment, we used only sH3N2 isolates with a collection date prior to the respective vaccine selection meeting for each season individually and calculated the genealogy and SD plots.

### Alignment & Phylogenetic inference

We generated multiple sequence alignments from both the nucleotide or amino acid sequences with MUSCLE^62^. To avoid shifts in the numbering of both alignments, positions with gaps in more than 80% of the sequences are removed from the alignment with TrimAl^63^. A phylogenetic tree was inferred from the nucleotide alignment using fasttree^64^ with the GTR-model, which enables a quick tree computation for large numbers of sequences while retaining a good accuracy compared to slower methods. For the pH1N1 influenza virus subtype, the A/California/05/2009 strain and for the sH3N2 influenza virus subtype the A/Moscow/10/1999 strain were used as an outgroup to root the trees. Both viruses were predominantly circulating in the viral population in the first season of our analyses^2,65^. We resolved multifurcations by adding further nodes and zero length edges into the outgoing edges to obtain a binary representation of the tree and subsequently applied the parsimony model of Fitch’s algorithm with accelerated transformation (ACCTRAN)^66^ to reconstruct amino acid sequences for the internal nodes of the phylogenetic trees. Amino acid changes were then inferred from the node-associated amino acid sequences and mapped to branches of the tree.

### Protein structure analyses

To investigate the structural properties of sweep-related changes, we mapped all sweep-related changes in HA1, NA, NP and the polymerase (PB1, PB2 and PA) onto the respective structures. For NA, the crystal structure of the A/California/07/2009(H1N1) strain (PDB 4B7Q) was used that covers residues 83-469 of NA. For HA1 and NP, we generated homology models based on structures from the RSCB database and the amino acid sequence of the A/California/07/2009(H1N1) strain using MODELLER^67,68^. Multiple sequence alignments containing protein templates and target sequences for modeling were calculated with MUSCLE^62^. Two structures (PDB 3M6S and 3LZG) from the A/Darwin/2001/2009(H1N1) strain displaying a sequence identity of 99.07% within residues 1-322 compared to the A/California/07/2009(H1N1) sequence were used as templates for composite modeling. For NP, we used the crystal structure of NP from the A/Wilson-Smith/1933(H1N1) strain (PDB 3RO5) with an identity of 91.9% compared to the A/California/07/2009(H1N1) strain. To obtain a three dimensional model of the viral polymerase, we generated individual models of PA, PB1 and PB2 for the A/California/05/2009(H1N1) strain using the Phyre II server^69^. These models were subsequently superposed onto the structure of the intact influenza A polymerase of *Reich et al.*^70^ (PDB 4WSB).

We determined the exposure of each site with the tool SURFACE from the CCP4 toolkit^71,72^ and calculated the accessible surface area (ASA), using a 1.4 Å radius for the probe sphere equal to the radius of water and the van der Waals radii for different types of atoms defined by Chothia^73^, as well as a step variable of 0.1. To calculate the relative solvent accessibility (RSA), we normalized ASA values with a theoretical value of the maximum possible solvent accessibility and classified residues as buried when the RSA is ≤ 0.05 and as exposed otherwise^74^.

## Data and Software Availability

The SD plots software, the figure and tables from this manuscript, and all related data used in this publication are fully available under: https://github.com/hzi-bifo/SDplots.

## Acknowledgements

This work was supported by the Helmholtz Centre for Infection research. The Heinrich Pette Institute, Leibniz Institute for Experimental Virology, is supported by the Free and Hanseatic City of Hamburg and the Federal Ministry of Health.

## Author Contributions Statement

A.C.M conceived the study. A.C.M and T.R.K planned and coordinated the study. A.C.M., S.R., T.R.K. and F.K. designed the methodology. T.R.K., S.R., J.L. and K.M. maintained the data, implemented the software and created results. A.C.M., S.R., J.L., T.R.K., G.G. and T.K. analyzed the results. T.R.K., S.R., J.L. and T.K. generated all visualizations. T.R.K., A.C.M. and S.R. wrote the original draft of this manuscript and all authors reviewed the manuscript.

